# Sustained attention under load: Neurophysiological mechanisms and behavioural consequences

**DOI:** 10.64898/2026.08.17.745232

**Authors:** Louise Catheryne Barne, Xuran Fu, Nilli Lavie

## Abstract

‘Sustained attention’ research demonstrates the negative consequences of declining task focus with time-on-task. Separate research demonstrates the critical role of perceptual load in task focus: increased perceptual load is shown to draw neural energy into task-relevant, rather than irrelevant, processing (Bruckmaier et al., 2020) thus improving task focus (Lavie, 2005). However, how perceptual load affects the neurophysiological mechanisms underlying sustained attention decline with time-on-task remains unknown.Moreover, research on the aperiodic component of EEG has linked the 1/f slope with excitation-inhibition (E/I) balance: steeper slopes reflecting reduced E/I ratio (Gao et al., 2017), but whether sustained attention decline can be attributed to a change in the aperiodic slope is an open question. Here we recorded EEG during gradual continuous-performance task (detecting infrequent mountain-among-city scenes), under low or high perceptual load (with or without overlaid ‘salt-and-pepper’ noise, respectively), and analysed periodic and aperiodic components. Time-on-task resulted in a widespread increase in alpha power and a steeper 1/f slope in a left temporoparietal cluster, accompanied by reduced detection sensitivity, increased response variability, and increased mind wandering, with reduced thought detail. Perceptual load improved task focus, indexed by reduced mind wandering, but exacerbated the effect of time-on-task on detection sensitivity, and the 1/f slope was steeper with time-on-task in a right parieto-occipital cluster under increased load. Overall, these findings suggest that sustained attention decline with time-on-task can be attributed to depletion of neural energy needed for excitatory signalling, which is further drained with increased processing demands in tasks of high perceptual load.

**Significance statement:** The ability to sustain attention on a task is known to rapidly decline over time, with costs ranging from ineffective performance at work or education to safety-critical consequences (e.g. in driving). This study identifies a novel neurophysiological mechanism underlying this decline by examining aperiodic electrophysiological activity (i.e. the 1/f slope of broadband power decline across the oscillation frequency spectrum) as a function of time-on-task and perceptual-processing demands (i.e. perceptual load). Our findings indicate that declining task performance with time-on-task was associated with a steeper 1/f slope, which is thought to reflect reduced excitatory over inhibitory neural activity. Higher perceptual load exacerbated both effects, suggesting that time-on-task drains neural resources required to sustain excitatory signalling, particularly under higher processing demands.

## Introduction

Limits on neural energy available for mental processing require focusing attention on task-relevant information. Although sustaining task-selective focus is critical for maintaining effective processing throughout the task period, ‘focused attention’ and ‘sustained attention’ have typically been studied in different bodies of research. Research of focused attention has established perceptual load as a major determinant of the ability to selectively focus on task-relevant information (e.g., Lavie, 1995; Lavie et al., 2014). Many studies have shown that tasks of increased perceptual load (e.g. greater perceptual processing complexity, or more subtle discrimination) result in both i) increased neural activity related to task-relevant processing, e.g. increased fMRI signal, evoked potentials, and alpha band (8-12 Hz) desynchronization, reflecting task-related increase in cortical excitability (Lavie, 2005; Molloy et al., 2015), and ii) reduced neural response to distractors, with increased distractor-related alpha power (Gutteling et al., 2022; Schwartz et al., 2005), and reduced distractibility from both external (Forster & Lavie, 2008; Lavie, 2010) and internal ‘mind wandering’ sources (Bruckmaier et al., 2023; Forster & Lavie, 2009).

Research of sustained attention has established a decline in attention with longer time-on-task (See et al., 1995; Warm et al., 2008), reflected in decreased accuracy, increased response variability, as well increased rates of mind wandering (Thomson et al., 2015; Krimsky et al., 2017; Martinez-Perez et al., 2023), and ‘out of the zone’ states (Esterman et al., 2013; Solís-Vivanco et al., 2024).

Time-on-task impact on sustained attention is known to involve increased alpha power (Bonnefond et al., 2011; Pershin et al., 2023), reflecting reduced task processing (e.g. Macdonald et al., 2011), and increased midfrontal theta power (Tran et al., 2020) though of reduced inter-trial consistency (Reteig et al., 2019). Midfrontal theta power is thought to indicate cognitive control, and its increase with time-on-task suggests increased demand on cognitive control due to increased attention lapses, evidenced for example by increased mind-wandering rates (Braboszcz & Delorme, 2011; Dias da Silva et al., 2022). These effects stress different accounts for the sustained attention decline with time-on-task, emphasizing either the increased challenge on cognitive control, or the reduced task-related cortical excitability.

However since the bulk of sustained attention research did not manipulate perceptual load, it remains unclear whether increased perceptual load would enhance the ability to sustain attention focus over time, by reducing neural processing of task-irrelevant distractors, and mind wandering, and hence reducing demand on cognitive control. Or whether instead the increased task-related neural processing demand, would drain neural energy that is required to sustain vigilant attention over time (Esterman & Rothlein, 2019).

We therefore investigated behavioural and neurophysiological measures of sustained attention under different levels of perceptual load in the present study. We measured both periodic (allowing examination of time-on-task effects on alpha and theta power) and aperiodic components of the EEG power spectrum. The aperiodic 1/f slope was of particular interest since it is thought to indicate the balance of excitatory versus inhibitory (E/I) activity (Gao et al., 2017): steeper slope (i.e. greater decline of broadband power with increased frequency) is taken to reflect decreased E/I ratio (Chini et al., 2022; McKeon et al., 2024) States of lower cortical excitability have accordingly been associated with steeper slopes, and reduced E/I ratio (Lendner et al., 2020; Waschke et al., 2021).We hypothesized that, in addition to the expected periodic effects, if sustained attention decline is mainly due to the draining of neural energy mediating task-related activity over time, a steeper 1/f slope would be expected with time-on-task (e.g., in parietal-occipital channels mediating task processing), and the further increase in neural energy demands with higher perceptual load would intensify the drainage of neural energy, hence would be expected to magnify the time-on-task-related steepening of the 1/f slope. In contrast, since increased perceptual load would reduce the draw on active cognitive control processes, it should reduce any time-on-task effect on the 1/f slope reflecting cognitive control modulations (e.g. in midfrontal channels).

## Materials and Methods

### Experiment 1

#### Participants

Thirty-four adult participants (30 females; average 23 years; age range 18-34; two left-handed) were recruited online via a UCL platform. The sample size was determined a priori to detect a within-participant effect of medium size (Cohen’s d ≥ 0.5, paired t-test) with 80% power. Participants had normal or corrected-to-normal vision and no reported history of neurological disorders. This study was approved by the UCL Research Ethics Committee. All participants provided written informed consent and were reimbursed £20 for their time.

#### Apparatus

Stimuli were generated in MATLAB (R2019a, MathWorks) using the Psychophysics Toolbox (Brainard, 1997) and presented on a Dell S2417DG monitor (resolution of 2560 x 1440 pixels, a refresh rate of 120 Hz, width of 52.7 cm). Eye data were recorded during the experiment using an Eyelink 1000 eye tracker (EyeLink, SR Research Ltd., Mississauga, Ontario, Canada) but not currently analysed. Participants had their head movements restrained with a chin rest located 63 cm from the screen.

EEG data were continuously recorded using the BioSemi Active Two system (BioSemi, Amsterdam, Netherlands), digitised at a 2048 Hz sample rate, with 24-bit A/D conversion, from 64 active scalp Ag/AgCl electrodes placed according to the international standard 10–10 system on a nylon head cap. Two extra bipolar electrodes were placed on the left and right mastoids for offline reference. Channels were adjusted to below the threshold of 40 microvolts offset, measuring impedance by the half-cell potential of the electrode interface before the start of the experiment. During the procedure, they were grounded to common-mode-sense (CMS) and driven-right-leg (DRL) electrodes. EEG analyses were performed using FieldTrip software (Oostenveld et al., 2011). Repeated measures ANOVA (including the Bayesian approach) were performed in JASP, Version 0.17.1.0 (JASP Team, 2018).

#### Stimuli and task procedure

Participants performed the gradual continuous performance task (gradCPT, Figure 1a) similar to Esterman et al. (2013), in which they were instructed to monitor 8-minute-long streams for infrequent scenes.

**Figure 1.**
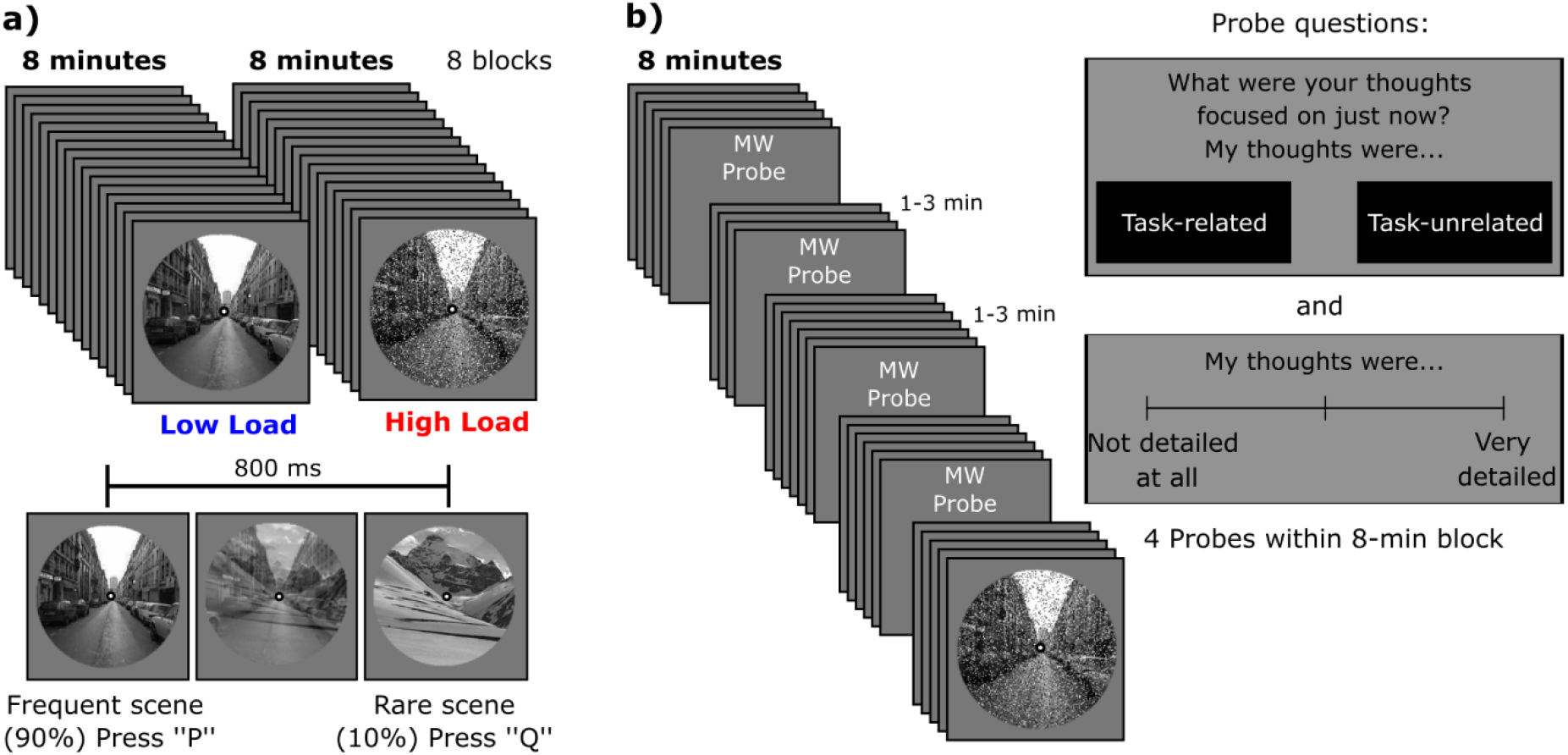
Task structure. a) Experiment 1. Participants had to monitor eight-minute-long streams of natural scenes with gradual onsets for frequent and infrequent scenes. High perceptual load blocks included an overlaid salt and pepper noise mask (19%) on the scenes;b) Experiment 2 had a similar structure to Experiment 1 but counted with mind wandering (MW) probes. There was one probe every one to three minutes; four were presented within a block.

Stimuli were circular greyscale photographs (4.58 degrees of visual angle) of mountain (n=10) and city (n=10) scenes with an overlaid central fixation point (0.15 degrees of visual angle) on a light grey background, all presented sequentially at the centre of the screen. The transition between sequential scenes was gradual by applying a linear pixel-by-pixel interpolation of the two images at each frame presentation. Each of these image composites (image N) composed of *x%* of the previous scene and 100% - *x%* of the next scene, starting with 100% of the previous scene and 0% of image N, which is termed the transition point. The stream started with a circular mask composing of a scrambled image of all scenes which then gradually transitioned to image N. The transition to image N from 0 to 100% occurred in 800 ms, a period considered image N trial. Therefore, each 8-minute stream had 600 trials. Stimuli were presented in random with the constraint that no identical scenes were repeated on consecutive trials. City scenes were presented on 90% of the trials, and mountain scenes on 10% of the trials. Participants were instructed to monitor scenes on the computer screen and indicate whether they transitioned to a city or a mountain scene by pressing the ‘P’ key with their right-hand index finger for a city scene and the ‘Q’ key with their left-hand index finger for a mountain scene, on the pc keyboard. Participants were unaware of the exact probability of scene types. However, they were told in the instructions that city images would occur more frequently, while encouraged to detect both scene categories to the best of their ability.

Perceptual load was manipulated by reducing the scene discriminability, with the addition of a salt and pepper noise mask (19% noise level) overlaid on each image in the high perceptual load streams (Gutteling et al., 2022; Yi et al., 2004). An instruction screen informed participants about the scene visibility condition for the following stream stating whether that block had clear/reduced visibility and presenting an exemplar of scene (with/without a noise mask). Participants performed eight blocks (each of an 8-minute-stream), four blocks under each perceptual load level, in an ABBAABBA or BAABBAAB order fashion, counterbalanced across participants. There was a 30 s break after each block and participants had to press a key to start a block when ready.

Before the experiment and EEG setup, participants were trained with four practice blocks (two for each load condition) each involving a 30 s stream, always starting with the low perceptual load condition. During practice, block sequence was alternated, no breaks were required between blocks, and performance feedback was provided after each practice block by showing participants’ percentage accuracy, for mountain and city image trials separately.

During the experiment, no accuracy feedback was given. However, participants received a warning message at the end of the block which asked them to pay attention and talk to the experimenter if they detected either less than 20% of the mountain scenes, or less than 50% of the city scenes. The experimenter then encouraged the participant to focus on the task, suggesting that they take a longer break if they felt too tired.

#### Behavioural analyses

Reaction times (RTs) were calculated based on the transition point to a new image, using Esterman et al. (2013) procedure. Prior to RT calculation an algorithm assigning responses to an image was run. First unambiguous responses, considered as the fastest response occurring between 70% coherence of image N and 40% coherence of image N+1 (corresponding to RT of 560 to 1120 ms) were assigned. Then, for the remaining images without an assigned response (ambiguous trials), the interval to consider image N RT was expanded to 40% coherence of image N up to 70% coherence of image N+1 (RT between 320 and 1360 ms) and the fastest response within this range was assigned to the corresponding image N. Accuracy was computed after determining the response-image correspondence.

Performance measures were estimated for the first (0-2 min) and last (6-8 min) periods of each block and were submitted to a 2 x 2 repeated measures ANOVA with perceptual load (low, high) and time-on-task (0-2 min, 6-8 min) as factors. We applied a signal detection analysis considering correct responses to the rare (10% occurrence) mountain images as hits, and a mountain-response to frequent (90%) city images as a false alarm. False alarm and hit rates were then used to calculate the non-parametric estimate of sensitivity index (*A’*) (J. Zhang & Mueller, 2005). RT was described by fitting an ex-Gaussian distribution over the correct responses to frequent stimuli. Three parameters were estimated: mean (µ, *mu*) and standard deviation (σ, *sigma*) of the normal component, and mean (τ, *tau*) of the exponential component. We chose to fit ex-Gaussian over the normal distribution to account for the established right skewness of RTs.

#### EEG analyses

##### Pre-processing

The continuous EEG data were referenced to the average of left and right mastoid channels. No offline filters were applied and the data was first segmented into long (8-min) blocks where a baseline (−200 to 0 ms, prior to the block onset) was applied, and the channel- and block-wise linear trends (detrend) were subtracted from the traces. Subsequently, the blocks were segmented into overlapping trials of two seconds locked to the start of each image presentation. Data were down-sampled to 512 Hz, and channels showing an unusually high variance (through visual inspection of distributions using *ft_rejectvisual*) were repaired by replacing them with the weighted average of their neighbour channels (number of interpolated channels: *median* = 0, *Q*_*0*.*75*_ = 1, *max* = 5). Independent component analysis (standard runica method) was run to identify and remove components related to eye movements. Between one and two components reflecting blinks and/or saccades were removed per participant (M = 1.21; one component for 27 participants, two for 7). Trials with an unusually high variance were excluded from further analyses using a semi-automatic procedure (*ft_rejectvisual* function, rejected trials: *M* = 7.32%, *SD* = 4.94%).

##### Power analyses

Power spectrum density (PSD) was estimated for frequencies ranging from 3 to 40 Hz by running a Fast Fourier Transform with a Hanning taper on each frequent image (city scene) trial, time-locked to image onset (0–800 ms). Only trials in which both the current trial and the immediately preceding trial were correctly detected were included in this analysis. Infrequent target (mountain scene) trials were not included to avoid contamination from target detection-related activity. PSD was computed for each trial and then averaged over trials according to their conditions. In addition, evoked PSD was also obtained by averaging trials according to their conditions in the time domain (a.k.a., ERP analyses) before spectral decomposition, thereby retaining only phase-locked activity. Induced (non-phase-locked) power was calculated by subtracting the evoked PSD from the total PSD (see supplementary material). Conditions were the combination of levels of perceptual load (low and high) with time-on-task (0-2 min and 6-8 min), yielding one PSD per condition, channel and participant. Evoked responses to abrupt events have been shown to alter estimates of aperiodic (1/f) parameters (Gyurkovics et al., 2022). In our data, however, parameter estimates were highly consistent whether derived from total PSD or induced PSD, indicating that our findings are unlikely to be driven by evoked activity.

The resulting neural power spectra were decomposed into the periodic (oscillatory peaks) and aperiodic (1/f-like) components by using a spectral parameterisation algorithm FOOOF (Donoghue et al., 2020) implemented in FieldTrip via code from the Brainstorm toolbox. Power spectra were parameterised across the frequency range 3 to 40 Hz, and the settings for the algorithm were: peak width limits = [0.5, 12]; maximum number of peaks = 3; minimum peak height = 1 dB; proximity threshold (or peak threshold) = 2.0; and aperiodic mode = ‘fixed’. This was applied for each channel, condition, and participant.

Periodic components were evaluated as the power spectra without the aperiodic signal (by setting the output as *fooof_peaks)*. For the aperiodic component, we analysed the estimated exponent, which quantifies the regression slope between power and frequency in log-log space, and offset for each channel and participant.

#### Statistics

##### Repeated measures ANOVA

Simple main effects analyses were performed in case of significant interaction between main effects. In case of non-significant findings, Bayesian analyses of effects using matched models were reported to support evidence for the alternative hypothesis (BF_10_).

##### Multidimensional EEG data

We used two-tailed cluster-based permutation tests based on paired t-scores (Maris & Oostenveld, 2007) to assess significant effect of time-on-task (6-8 min - 0-2 min), perceptual load (High - Low), and interaction (effect of time-on-task in High - effect of time-on-task in Low) on the 1/f slope, 1/f offset, and oscillations. Tests were run with 2000 random permutations, channel neighbours (defined as in the FieldTrip neighbour layout template for the BioSemi 64-electrode cap) with a minimum number of two channels required to be included in the clustering (*minnbchan* = 2), cluster statistics as the maximum of the summation, and cluster alpha of 0.05.

We computed Cohen’s d across all channels and, where applicable, all frequencies, and report the maximum value as the largest estimated effect size within our EEG dataset. For clusters showing significant differences in the cluster-based tests, we additionally report Cohen’s d of the signal averaged over the spatial (and, where applicable, spectral) extent of the cluster, providing a more conservative effect size estimate.

### Experiment 2

#### Participants

Twenty-four new participants (19 females, *M* = 24 years, age range 19-46; one left-handed) completed Experiment 2. We adopted this sample size to detect a within-participant effect (Cohen’s d ≥ 0.6, paired t-test) with 80% power based on the time-on-task effects in Experiment 1. Participants were recruited via a UCL participant pool platform using the same criteria as Experiment 1. All participants provided written informed consent and were reimbursed £15 for their time.

#### Stimuli and task procedure

The experiment was run on a Dell P2417H monitor (1920 x 1080 resolution, 60 Hz refresh rate). Viewing distance was 63 cm from the screen, fixed with a chinrest. Stimuli and procedure were the same as in Experiment 1, except for the following changes. The stimuli were 6.11 degrees of visual angle (slightly larger than in Experiment 1), and thought probes were added to assess whether participants’ thoughts during the task were task-related or task-unrelated, as well as the thoughts’ level of detail (Figure 1b). Examples of thoughts were given during the instructions, for task-unrelated thoughts (TuTs) these were ‘What will I have for dinner?’ or ‘I need to contact my friend after this experiment’. For task-related thoughts these were ‘When will the next target appear?’ or ‘Oops! I pressed the wrong button.’ Participants were requested to respond honestly, and informed that there was no penalty to reporting task-related or task-unrelated thoughts.

The thought probes followed each other on separate screens, to which participants answered by moving and clicking the mouse with their right hand. The first question, ‘What were your thoughts focused on just now?’ required a two-alternative forced choice response, by clicking one of the response boxes ‘task related’ or ‘task-unrelated’ that followed the statement “my thoughts were”.

The second question asked whether those thoughts were detailed and clear. A slider going from ‘not detailed at all’ to ‘very detailed’ was shown, and participants responded by clicking at the position matching their experience. The middle point could not be clicked. After they had answered the two questions, they pressed ‘P’ on the keyboard with their right hand to continue the task.

One thought probe was inserted in the middle (at 15 s) of each 30 s long practice block. In the main experiment, thought probes were presented four times within each 8-minute block stream, in pseudo-random intervals ranging from one to three minutes. Thus, the first probe had to be presented at any point between the 1^st^ and 3^rd^ minute of the stream and the fourth probe at any moment from the 5^th^ to 7^th^ minute. The specific pseudo-random intervals assigned were repeated across a pair of blocks of an even number to ensure equal sampling time across perceptual load conditions. Additionally, no infrequent stimuli (mountain scenes) were presented during the last 12 s preceding a thought probe (since these can in principle trigger task-related thought). The experimental session (including the training) lasted approximately one hour and 40 minutes.

#### Analyses

Blocks where participants detected less than 50% of city images (i.e., false alarm rate of 50% or higher) were excluded from all analyses (this resulted in the exclusion of one block for one of the participants). The behavioural analyses of the effect of perceptual load on sustained attention were the same as in Experiment 1.

To assess the effects of perceptual load and time-on-task on TuTs and their level of detail, we ran mixed effects analyses using JASP, Version 0.17.1.0 (JASP Team, 2018). The likelihood ratio test (F-test, Type III Sum of Squares) was applied to compute degrees of freedom for p-values. Time-on-task in these models was based on the probe order (ordinal variable expressing the probe sampling order within the block, ranging from 1 to 4).

Aa generalised linear mixed effects model (Binomial distribution with Cauchit link function) was run with the fixed effects of perceptual load, time-on-task, and their interaction to predict the occurrence of TuTs. TuT occurrence was modelled as the number of “successes” when asking such a yes-no question a few times. We included intercepts for each participant as a random effect in the model.

A linear mixed effects model (Gaussian distribution with Identity link function) was run with the same fixed and random effects, on the level of TuTs’ detail (a continuous variable from 0 to 100% detail).

## Results

### Experiment 1

In Experiment 1, participants performed the gradCPT while EEG was recorded. We first examined whether perceptual load and time-on-task affected response distribution and detection sensitivity.

The mean RT (*mu*, Figure 2a) was increased with increased perceptual load (*F*_(1,33)_ = 93.12, *p* < .001, 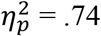) and with longer time-on-task (*F*_(1,33)_ = 21.93, *p* < .001, 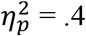). These results confirmed the increased demand on attention both with perceptual load and with time-on-task. Figure 2a also shows a trend for an interaction indicating larger effect of time-on-task in conditions of high (vs. low) perceptual load, which failed to reach significance (*F*_(1,33)_ = 3.47, *p* = .071, 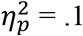), and *BF*_*10*_ = 0.96, suggested anecdotal evidence for H_0_.

**Figure 2.**
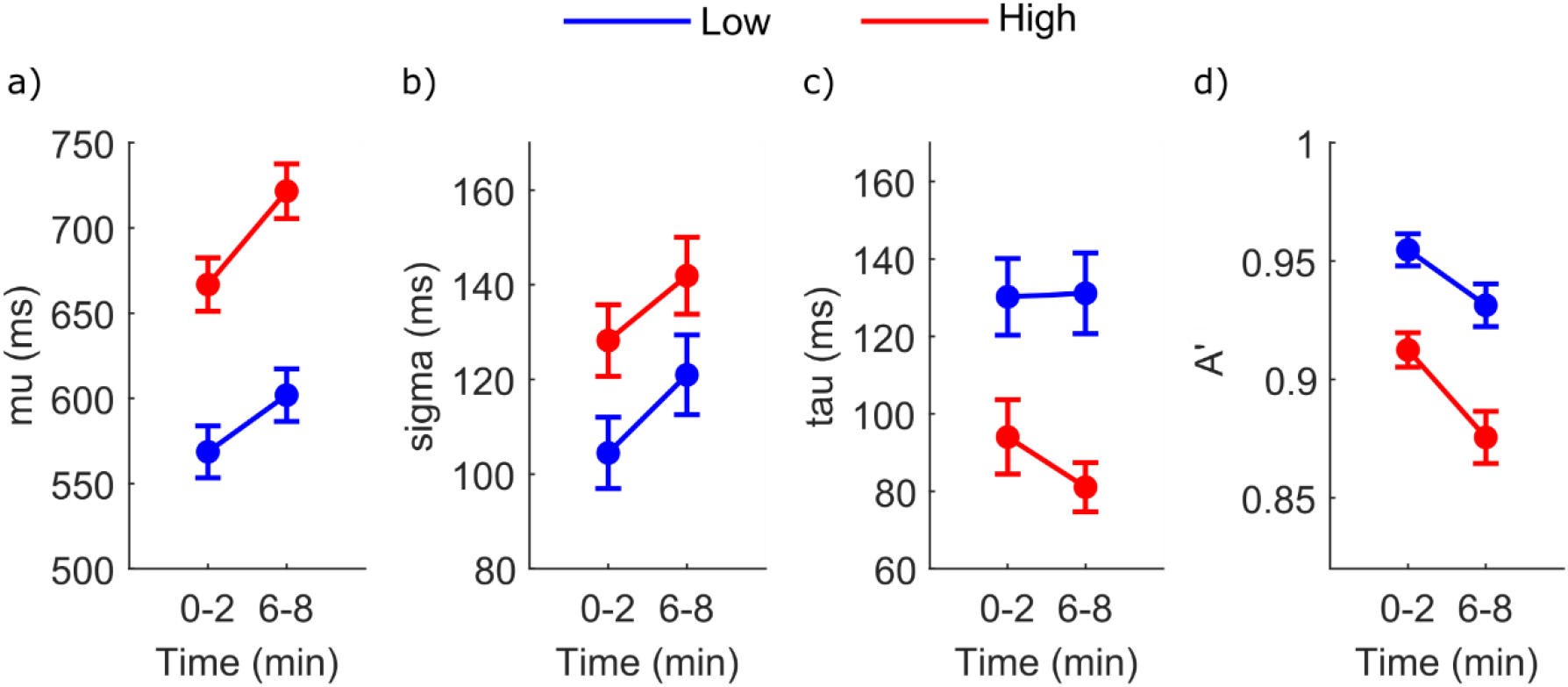
Performance as a function of time-on-task (0-2, 6-8 min) and perceptual load (low, high) in experiment 1. Error bar represents the standard error (SE; mean ± 1 SE). a) RT mean (mu); b) RT variability (sigma); c) RT tail (tau); d) Detection sensitivity (A’).

Higher demands on attention were additionally corroborated with increased RT variability (*sigma*, Figure 2b) with both higher perceptual load (*F*_(1,33)_ = 29.63, *p* < .001, 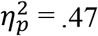) and longer time-on-task (*F*_(1,33)_ = 15.34, *p* < .001, 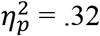). The interaction was not significant (*F*_(1,33)_ = 0.28, *p* = .601, 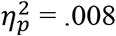), and *BF*_*10*_ = 0.31, indicated moderate evidence for H_0_.

The tail of RT distribution (*tau*, Figure 2c) was reduced under high compared to low perceptual load (*F*_(1,33)_ = 35.92, *p* < .001, 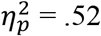). There was no significant effect of time-on-task (*F*_(1,33)_ = 0.68, *p* = .42, 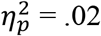, with *BF*_*10*_ = 0.32 showing moderate evidence for H_0_) and no interaction (*F*_(1,33)_ = 1.7, *p* = .202, 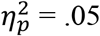, *BF*_*10*_ = 0.57 only suggesting anecdotal evidence for H_0_). Corroborating previous findings (Yamashita et al., 2021), these results supported that failures of attention are better expressed in RT by increases in *mu* and *sigma* rather than in *tau* when performing rhythmic tasks like the gradCPT.

Detection sensitivity (*A’*, Figure 2d) was reduced with increased perceptual load (*F*_(1,33)_ = 74.21, *p* < .001, 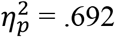) and increased time-on-task (*F*_(1,33)_ = 28.89, *p* < .001,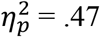). Moreover, the impairment in detection sensitivity with time-on-task was greater under a higher level of perceptual load (interaction: *F*_(1,33)_ = 4.14, *p* = .05, 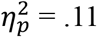; simple main effect of time-on-task under low load: *F* = 13.08, *p* < .001, Cohen’s *d* = 0.62; simple main effect of time-on-task under high load: *F* = 31.63, *p* < .001, Cohen’s *d* = 0.96). This result suggested that task-related information processing was compromised over prolonged periods, especially when the task involved increased demands on perceptual processing.

#### Broadband aperiodic power

We proceeded to EEG analyses by using the *fitting oscillations and one-over-f* algorithm (Donoghue et al., 2020) to dissociate oscillations from the aperiodic 1/f signal at the level of each condition, electrode, and participant on the total PSDThe goodness-of-fit measured by *R*^*2*^ *Mdn* Low _0-2 min_ = .99 (*IQR* = .96, 1); *Mdn* High _0-2 min_ = .98 (*IQR* = .95, .99); *Mdn* Low _6-8 min_ = .99 (*IQR* = .96, .99), *Mdn* High _6-8 min_ = .99 (*IQR* = .96, .99), supported reliable estimates for inspecting the parameters of the aperiodic signal (slope and offset) and the power of oscillations (i.e., the power spectrum with the 1/f aperiodic signal removed).

The cluster-permutation test revealed a significant effect of time-on-task on the 1/f slope (*p* = .038, Cohen’s *d* of circumscribed cluster = 0.61, and maximum Cohen’s *d* of 0.55 over T7, Figure 3a), reflecting steeper slope with longer time-on-task.

**Figure 3.**
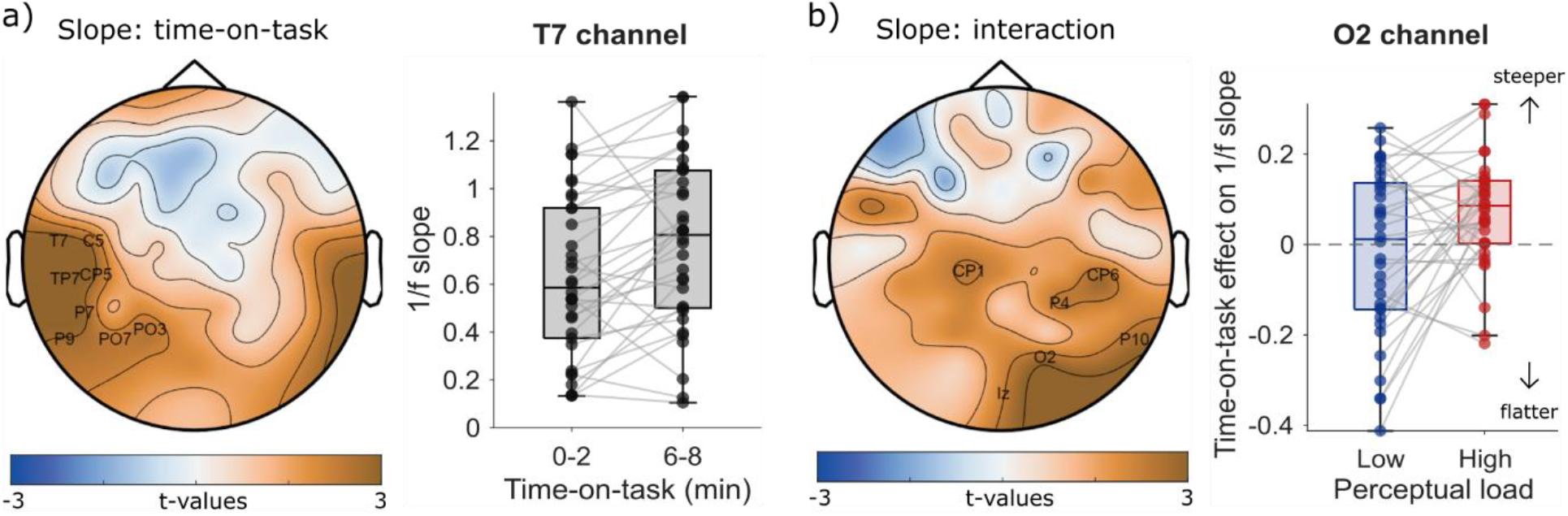
1/f slope. a) Right: scalp topography of the main effect of time-on-task (6-8 min minus 0-2 min), channel labels are shown for the significant cluster. Left: Boxplot of the 1/f slope in channel T7 as a function of time-on-task. Individual participant data are overlaid and connected across conditions. b) Scalp topography of the interaction of time-on-task with perceptual load (time-on-task effect on high load minus time-on-task effect on low load). Labels are shown for the significant cluster of channels. Left: Boxplot of slope differences with time-on-task (6–8 min minus 0–2 min) at channel O2 under different levels of perceptual load. Positive values represent an increase in slope over time-on-task. Individual participant data are overlaid and connected between perceptual load conditions.

An interaction of load and time-on-task was also found (*p* = .048, Cohen’s *d* of circumscribed cluster = 0.55, and maximum Cohen’s *d* of 0.48 over O2, Figure 3b). As shown in Fig 3b this interaction reflected that time-on-task resulted in a larger effect on the 1/f slope in the high load than the low load condition. This was also confirmed in a simple main effect analysis on the averaged exponent over the channels that composed the right-lateralized interaction cluster (CP1, CP6, P4, P10, O2, and Iz). The 1/f slope was steeper with longer periods on the task of high perceptual load (0-2 min: M = 1.05, SE = 0.05, 6-8 min: M = 1.13, SE = 0.04, *F* = 10.98, *p* = .002, Cohen’s *d* = 0.57). However, there was moderate evidence for no change in the slope as a function of time-on-task in low perceptual load (0-2 min: M = 1.09, SE = 0.05, 6-8 min: M = 1.09, SE = 0.05, *F* = 0.02, *p* = .9, Cohen’s *d* = 0.02, BF_10_ = 0.185). Finally, no cluster was found for the main effect of perceptual load.

For the 1/f offset, cluster-permutation test revealed a significant effect of time-on-task (p = .002, Cohen’s *d* of circumscribed cluster = 0.74, and maximum Cohen’s *d* of 0.89 over T7, Figure 4). The interaction cluster did not reach statistical significance (p = .066) and no cluster was found when testing for an effect of perceptual load.

**Figure 4.**
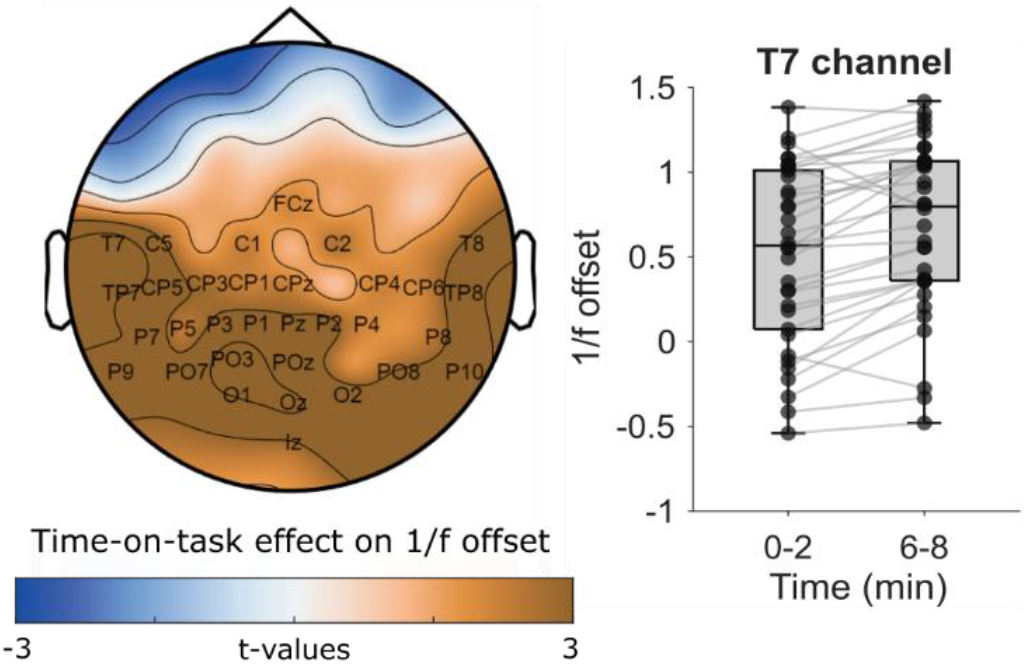
1/f offset. Right: scalp topography of the main effect of time-on-task (6-8 min minus 0-2 min), channel labels are shown for the significant cluster. Left: Boxplots of the average 1/f offset over channel T7 as a function of time-on-task. Individual participant data are overlaid and connected across conditions.

#### Periodic components

The analysis of oscillations showed that the power of frequencies ranging from 6 to 14 Hz (peak over 10 Hz) was enhanced with longer time-on-task, which was evident over all channels (*p* < .001, Cohen’s *d* of circumscribed cluster = 1.02, and maximum Cohen’s *d* of 1.21 over P10 at 10 Hz, Figure 5). The effect broadens the conventional alpha band range (8-12 Hz), but peaking at 10 Hz and gradually diminishing away from this frequency. There were no significant clusters for the effect of perceptual load (*p* = .416) or the interaction (*p* = .524).

**Figure 5.**
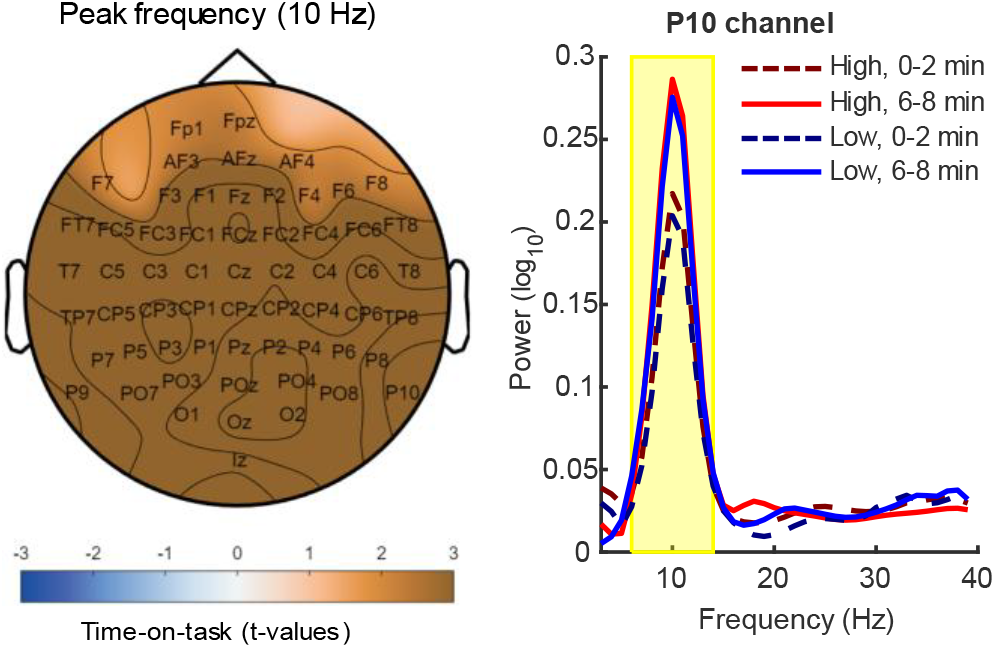
Periodic component. Right: Scalp topography of time-on-task effect at 10 Hz (6-8 min minus 0-2 min), channel labels are shown for significant cluster of channels. Left: Average power spectrum (oscillatory fit with 1/f regressed out) over participants at channel P10 across time-on-task and perceptual load conditions. Yellow shadow represents the significant frequency range (6-14 Hz) for time-on-task effect.

These EEG markers together with behavioural results suggest that perceptual load increased sustained attention failures with longer time-on-task due to a decrement in the ability to sustain higher-demands on excitatory neural transmission (in high vs. low perceptual load) over time. Alternatively, one may argue that the high perceptual load task induced greater ‘task-fatigue’, resulting in a failure of cognitive control to prioritize task-related information, over task-unrelated thoughts, which are by nature more related to a person’s interests, with longer time-on-task. Thus, a failure of cognitive control is expected to lead to a greater increase in mind wandering with time-on-task in the high compared to low perceptual load conditions. In contrast, if perceptual load did draw more resources into the task relevant processing, rather than reducing the ability to exert cognitive control with longer time-on-task, we would expect that increased perceptual load would reduce the occurrence of task-unrelated processes mediating mind wandering.

To tease apart these accounts we tested the effects of perceptual load on task-unrelated thoughts (mind wandering) as a function of time-on-task in Experiment 2.

### Experiment 2

In this experiment the gradCPT included mind wandering (MW) probes randomly interspersed among the task trials, requiring participants to report whether their thoughts were task-related or task-unrelated (i.e. MW) and to indicate the level of thought detail on an analogue scale.

Performance measures of RT and detection sensitivity replicated Experiment 1’s results. As can be seen in Figure 6a, responses (Mean RT, *µ*) were slower with higher perceptual load (*F*_(1, 23)_ = 49.12, *p* < .001, 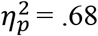), and slower over time-on-task (*F*_(1, 23)_ = 15.09, *p* < .001, 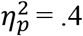). The interaction of load and time-on-task was not significant (*F*_(1, 23)_ = 2.38, *p* = .137, 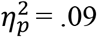 *BF*_*10*_ = 0.78, anecdotal evidence for H_0_).

**Figure 6.**
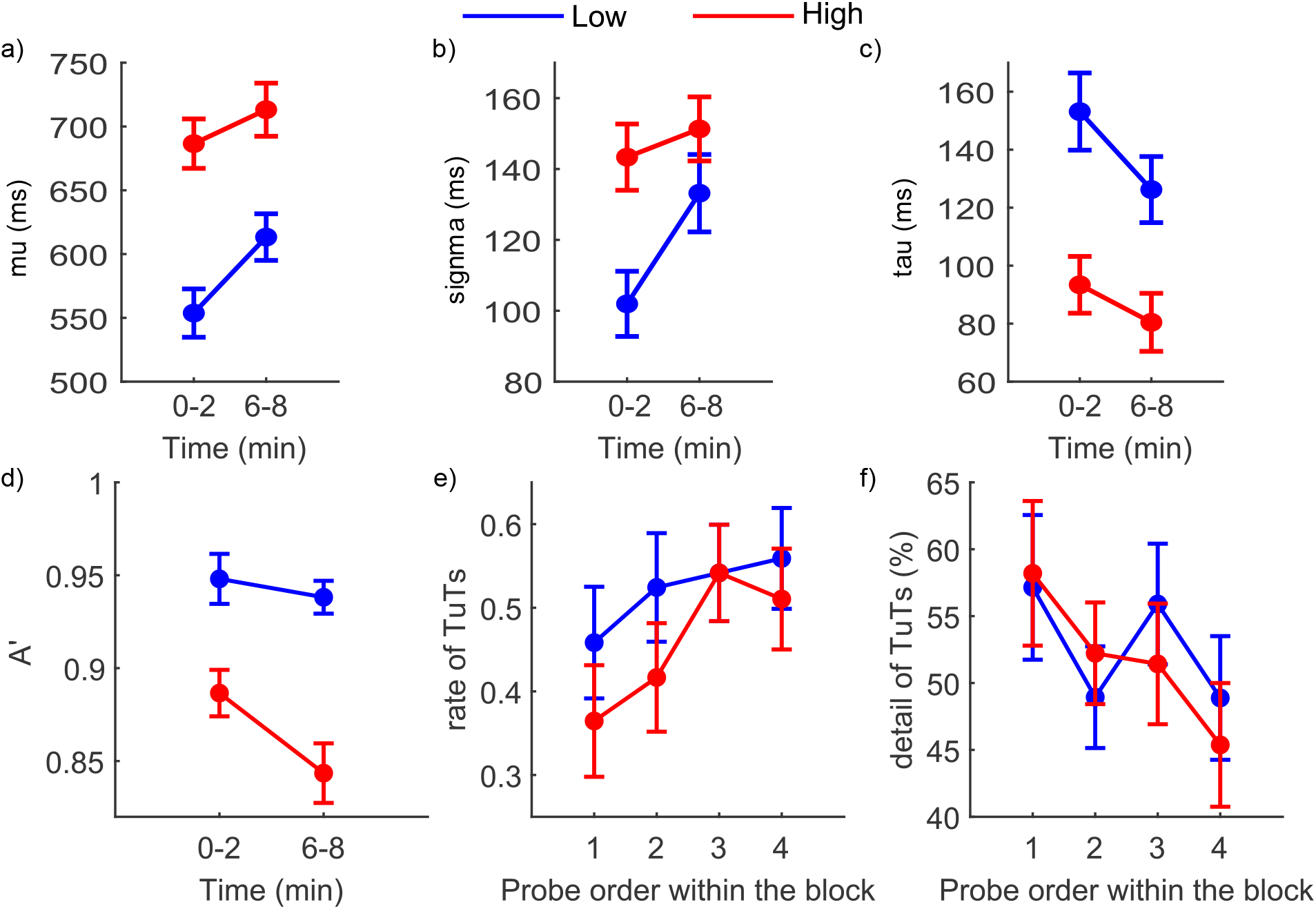
Performance as a function of time-on-task and low (in blue) and high (in red) perceptual load in Experiment 2. Error bars represent the standard error (SE; mean ± 1 SE). a) RT mean (mu); b) RT variability (sigma); c) RT tail (tau); d) Detection sensitivity (A’); e) Occurrence rate of task-unrelated thoughts (TuTs); f) Level of detail of TuTs.

Variability of responses (Figure 6b; σ) was also higher with time-on-task (*F*_(1, 23)_ = 8.26, *p* = .009, 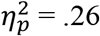) and perceptual load (*F*_(1, 23)_ = 20.33, *p* < .001, 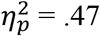). However their interaction was not significant (*F*_(1, 23)_ = 3.09, *p* = .092, 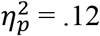; *BF*_*10*_ = 1.52, anecdotal evidence for H_1_).

Analysing the exponential tail of RT distributions (Figure 6c; *τ*), we found that high perceptual load reduced the tail compared to low perceptual load, as before (*F*_(1, 23)_ = 27.90, *p* < .001, 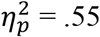). Time-on-task decreased skewness of RT distribution (*F*_(1, 23)_ = 4.84, *p* = .038, 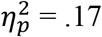), with no interaction (*F*_(1, 23)_ = 0.72, *p* = .404, 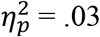; *BF*_*10*_ = 0.4, anecdotal evidence for H_0_). Repeated measures ANOVA of detection sensitivity revealed a main effect of perceptual load (*F*_(1, 23)_ = 83.38, *p* < .001, 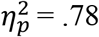) and of time-on-task (*F*_(1, 23)_ = 15.77, *p* < .001, 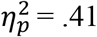), as can be seen in Fig 6d detection sensitivity was reduced with both higher perceptual load and longer time-on-task. There was also an interaction (*F*_(1, 23)_ = 4.50, *p* = .045, 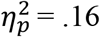), which reflected a larger effect of time-on-task in the high load (simple main effect of time: *F* = 15.43, *p* < .001, Cohen’s *d =* 0.8) than in the low load condition (where the simple main effect of time was not significant: *F* = 1.08, *p* = .310, Cohen’s *d* = 0.21, although *BF*_*10*_ = 0.35, only provided anecdotal evidence for H_0_). This pattern mirrored the effects of perceptual load and time-on-task on the aperiodic 1/f slope.

Perceptual load also reduced the rates of mind wandering (see Figure 6e): perceptual load was a significant predictor of the occurrence of mind wandering in the GLME model (χ^2^(1) = 4.72, *p* = .030), with lower probability rates estimated in high load (*Estimate of marginal means (EMM)* = .43, *95% CI* = [.33, .55]) compared to low load (*EMM* = .53, [.41,.64]). The model also indicated time-on-task (as in probe order) as a significant factor (χ^2^(3) = 9.78, *p* = .021), with the likelihood of TuT reports increasing progressively with longer periods on task (1: *EMM* = .39, [.29, .52]; 2: *EMM* = .45, [.33, .58]; 3: *EMM* = .55, [.42, .67]; 4: *EMM* = .54, [.41, .66])), (Figure 6e). Together with the evidence of time-related decline in the task performance, this result indicated a deterioration in the ability to control the task focus with longer periods on task. Lastly, the interaction between load and time-on-task was not significant in predicting mind wandering occurrence (χ^2^(3) = 1.89, *p* = .595).

Time-on-task (probe order) was a significant factor in predicting the level of thoughts detail (χ^2^(3) = 8.12, *p* = .044; Figure 6f), with less detailed thoughts estimated over time (1: *EMM* = 58, [50.31, 65.6]; 2: *EMM* = 49.75, [42.43, 57.08]; 3: *EMM* = 51.27, [44.19, 58.34]; 4: *EMM* = 47.51, [40.4, 54.62])). Perceptual load was not significant predictor (χ^2^(1) = 0.07, *p* = .792) and neither was the interaction of perceptual load and time-on-task (χ^2^(3) = 2.22, *p* = .528).

Thus, mind wandering increased but comprised less detailed thoughts as a function of time-on-task. Importantly, high perceptual load reduced the occurrence of task-unrelated thoughts compared to low perceptual load, demonstrating that high load improved task focus over the whole gradCPT period.

## Discussion

The present findings established that a decline in sustained attention over time involves a steeper 1/f slope of the EEG signal, with the largest effect at left temporo-parietal channels, and enhanced activity in the alpha frequency band, as well as impaired detection sensitivity, and slower and more variable RT. High perceptual load also reduced detection sensitivity in the task, as expected, together with slower and more variable responses.

Importantly, the results established that increased perceptual load results in a larger effect of time-on-task on the 1/f slope in parieto-occipital channels: this was steeper with longer time-on-task in conditions of high, compared to low, perceptual load. The decline in detection sensitivity with time-on-task was also larger in high (vs. low) perceptual load. Finally, MW rates increased and the level of thought detail decreased with time-on-task, however perceptual load reduced MW rates throughout the task period. These findings provide a new insight into the neuro-cognitive mechanisms of sustained attention and how these are impacted by perceptual load, as we discuss next.

### Effects of time-on-task and perceptual load on sustained attention

Our findings indicate that sustained attention over a longer period involves not only impairment in task performance and increased task-unrelated thoughts of less detail, but also increased alpha synchrony and a steeper 1/f slope. The EEG findings are thought to reflect reduced cortical excitability (Klimesch, 2012), and reduced excitatory over inhibitory neural signalling respectively (e.g. Gao et al., 2017; McKeon et al., 2024). Together with previous findings that a steeper 1/f slope is characteristic of rest compared to task (He et al., 2010); and of states of lower arousal (Lendner et al., 2020; Waschke et al., 2021) that result in states of lower consciousness (Maschke et al., 2023) we suggest that the effects of time-on-task on sustained attention involve not just reduced cognitive control (leading to increased MW) but also lower neural capacity to respond to external stimuli (as shown by the decrement in detection sensitivity), and to support more detailed thoughts.

Steeper 1/f slope is associated with less effortful listening (Woods et al., 2024) and reduced attention to the task-irrelevant modality in a multisensory detection task (Waschke et al., 2021), in line with our findings that reduced ability to sustain attention with time-on-task results in a steeper 1/f slope too.

Oddball stimuli, informative visual cues, as well as incongruent distractors are also associated with a steeper 1/f slope (Gyurkovics et al., 2022; Kałamała et al., 2024; Zhang et al., 2023), and these may arguably demand more attention. However, the stimulus-induced effects on the 1/f slope may reflect a transient state that interrupts ongoing processing, and trigger the update of neural representations, involving inhibition of the pre-stimulus activity. The former two studies as well as the present gradCPT paradigm employed continuous streams, in which any stimulus-onset effect on the spectral slope is minimised.

Importantly, the steepening of the 1/f slope with time-on-task was magnified under high perceptual load: a condition shown to increase the neural energy levels (as indicated by intracellular neural metabolic enzyme marker) related to attended processing (Bruckmaier et al., 2020). Thus, our results suggest that high perceptual load intensifies the drain on neural processing related to perception of the task stimuli with time-on-task, resulting in a greater deterioration of sustained attention with time, as reflected also in the greater time-on-task related decline in detection sensitivity with increased perceptual load.

Moreover, although time-on-task was associated with increased MW, perceptual load enhanced task focus throughout the task period and the increase in MW with time-on-task involved thoughts of reduced detail. Thus with reduced E/I in high perceptual load, the loss of cognitive control with time-on-task (which leads to increased MW) is moderated by reduced availability of neural resources to support detailed thoughts, since less-detailed thoughts are recognised as requiring less neural energy than detailed task-unrelated thoughts (Bruckmaier et al., 2023).

Our results also suggest that the main effect of time-on-task, and its interaction with perceptual load, were captured by distinct sets of channels, with the interaction effect more robustly expressed over right-lateralized posterior electrodes and the main effect over left temporal-parietal channels. This pattern suggests that there may be separable sustained attention functions involved in the task, as follows. The right lateralised system that is depicted by occipital-parietal channels is well known to be involved in sustained attention and vigilance over time (Hemmerich et al., 2025; Langner & Eickhoff, 2013) and our findings suggest that perceptual load exacerbates the time-on-task related decline in sustaining excitatory over inhibitory signalling in this system. In contrast, the main effect of time-on-task in a left temporal temporoparietal cluster may reflect an element of sustained attention, that declines with time irrespective of the task demand. For example, the maintenance of the mountain scenes template to monitor for, is a top-down function that may deteriorate with time, but unlikely to be affected by the level of perceptual load of the presented stimuli.

### 1/f slope and neural noise

Previous studies of ageing (Voytek et al., 2015) and ADHD (Pertermann et al., 2019) have interpreted a steeper 1/f slope as indicative of reduced neural noise, due to stronger autocorrelation of neural activity (e.g., synchronous spikes), as described by the Wiener-Khinchin theorem. According to this theorem, the signal retains a longer memory of its past when low frequencies dominate the spectrum. Thus, a steeper 1/f slope can also be interpreted as a marker of increased redundancy, reflecting less dynamic and more predictable (i.e., highly autocorrelated) signals. Such redundancy is crucial for error detection and correction especially when communicating over noisy channels (Shannon, 1948). In this study, we increased perceptual load specifically by adding noise to the visual images.Therefore, if prolonged periods of sustained attention drain processing capacity, the observed steeper 1/f slope may reflect a shift towards more redundant and autocorrelated neural coding, rather than increased information processing, particularly under the noisy condition induced by our perceptual load manipulation.

### Alpha oscillations

Time-on-task was associated with increased alpha power across all EEG channels. This provides support for time-on-task reducing cortical excitability, consistent with previous research (Pershin et al., 2023; Tran et al., 2020). However, we note that attention-related changes in both alpha power and EEG spectral exponents may arise from distinct neural mechanisms (Kosciessa et al., 2021; Waschke et al., 2021) and despite their correlation they cannot be reduced to each other (Cunningham et al., 2023; Manyukhina et al., 2024). Additionally, our findings indicate different scalp topographies for these two markers: time-related changes in alpha power are observed across all areas of the scalp, likely signifying reduced overall excitability. In contrast, changes in the aperiodic slope are more pronounced in the occipital parietal channels. Such topography may suggest that the aperiodic slope change is more pronounced within a specific task-related network that supports attention and perceptual capacity (Beck et al., 2001, 2006; Malhotra et al., 2009; Wojciulik & Kanwisher, 1999). Moreover, the time-on-task effect on 1/f slope was magnified in conditions of high perceptual load, known to increase the attentional demand, thus supporting an attention-draining account, the time-on-task effect on alpha power did not vary with perceptual load. Together with the generalised scalp activity, the effects of time-on-task on alpha power may then reflect a general effect of overall reduced cortical excitability in the course of the task period.

### 1/f offset

The 1/f offset was also found to increase with time-on-task in our study. The offset of the aperiodic signal has been shown to positively correlate with population spiking (Manning et al., 2009) and BOLD signal from fMRI (Winawer et al., 2013), and is thought to be predominantly affected by low frequencies (Frelih et al., 2025). Additionally, computational models suggest that lower frequencies tend to have more power when more ion channels are active in a network, regardless of whether these channels generate excitatory or inhibitory postsynaptic potentials (Gao et al., 2017). Therefore, while the aperiodic slope may reflect the balance between excitation and inhibition (E/I), a larger offset would likely reflect more overall network activity, but it is unclear whether this reflects greater inhibitory activity. Future research, particularly focused on LFP/EEG signals, is needed to further explore other potential explanations regarding 1/f offset.

### Conclusions

The present findings demonstrate that the decline in sustained attention with longer time-on-task (as manifested in impaired performance) is worsened with increased perceptual load in the task, despite its improvement of task focus, reflected in reduced mind wandering throughout the high-load task. A neurophysiological account for these results is offered by our findings of the interactive effect of time-on-task and perceptual load on the 1/f spectral slope: longer time-on-task was associated with a steeper 1/f slope, and increased perceptual load in the task amplified this effect.

Together with our findings of increased alpha power with time-on-task, and the view that a steeper slope reflects reduced excitatory relative to inhibitory neural activity, our findings suggest that time-on-task drains the neural resources required to sustain excitatory signalling, an effect further exacerbated by the increased neural energy demands of high perceptual load in the task.

## Supporting information

Supplementary analyses evaluating evoked responses and induced power power spectrum density parametrization.

## Acknowledgments

This work was supported by Toyota Motor Europe.

