## Supplementary analyses evaluating evoked responses and induced power power spectrum density parametrization. for "Sustained attention under load: Neurophysiological mechanisms and behavioural consequences"

**Supplemental Material**

Parametrization of induced PSD

The goodness-of-fit measured by $R_2$ Mdn Low 0-2 min = .99 (IQR = .96, .99); Mdn High 0-2 min = .98 (IQR = .95, .99); Mdn Low 6-8 min = .99 (IQR = .96, .99), Mdn High 6-8 min = .99 (IQR = .96, .99). Supplementary Figure 1 shows the scalp topography of time-on-task effects, and of their interaction with perceptual load, on the parameters of the induced power spectrum. Supplementary Figure 2 compares the evoked and total power spectra, showing the relative contribution of phase-locked activity to the total spectrum.

*1/f slope*

The cluster-permutation test revealed a significant effect of time-on-task on the 1/f slope of induced PSD (p = .034, Cohen's d of circumscribed cluster = 0.61, and maximum Cohen's d of 0.54 over TP7), reflecting steeper slope with longer time-on-task. An interaction of load and time-on-task was also found (p = .038, Cohen's d of circumscribed cluster = 0.61, and maximum Cohen's d of 0.49 over CP6, P10 and O2). The interaction was confirmed in a simple main effect analysis on the averaged exponent over the channels that composed the right-lateralized interaction cluster (FT7, FC5, CP1, P1, Iz, CP6, CP4, P10, and O2). The 1/f slope was steeper with longer periods on the task of high perceptual load (0-2 min: M = 1.02, SE = 0.05, 6-8 min: M = 1.10, SE = 0.04, F = 11.09, p = .002, Cohen's d = 0.57). However, there was moderate evidence for no change in the slope as a function of time-on-task in low perceptual load (0-2 min: M = 1.06, SE = 0.05, 6-8 min: M = 1.05, SE = 0.05, F = 0.09, p = .771, Cohen's d = -0.05, BF10 = 0.19). Finally, no cluster was found for the main effect of perceptual load.

*1/f offset*

For the 1/f offset of induced PSD, cluster-permutation test revealed a significant effect of time-on-task (p = .002, Cohen's d of circumscribed cluster = 0.75, and maximum Cohen's d of 0.87 over T7). There was a significant cluster found for the interaction (p = .030; Cohen's d of circumscribed cluster = 0.64, and maximum Cohen's d of 0.48 over O2). This was also confirmed in a simple main effect analysis on the averaged offset over the channels that composed the right-lateralized interaction cluster (CP1, P1, FT8, FC4, CP6, CP4, P10, and O2). The 1/f offset was higher with longer periods on the task of high perceptual load (0-2 min: M = 1.09, SE = 0.06, 6-8 min: M = 1.19, SE = 0.05, F = 27.44, p < .001, Cohen's d = 0.9). However, there was moderate evidence for no change in the offset as a function of time-on-task in low perceptual load (0-2 min: M = 1.12, SE = 0.05, 6-8 min: M = 1.14, SE = 0.05, F = 0.98, p = .330, Cohen's d = 0.17, BF10 = 0.29). No cluster was found when testing for an effect of perceptual load.

*Periodic component*

The analysis of oscillations of induced PSD showed that the power of frequencies ranging from 6 to 14 Hz (peak over 10 Hz) was enhanced with longer time-on-task, which was evident over all channels ($p < .001$, Cohen's d of circumscribed cluster = 1.01, and maximum Cohen's d of 1.20 over P10 at 10 Hz). There were no significant clusters for the effect of perceptual load or the interaction.

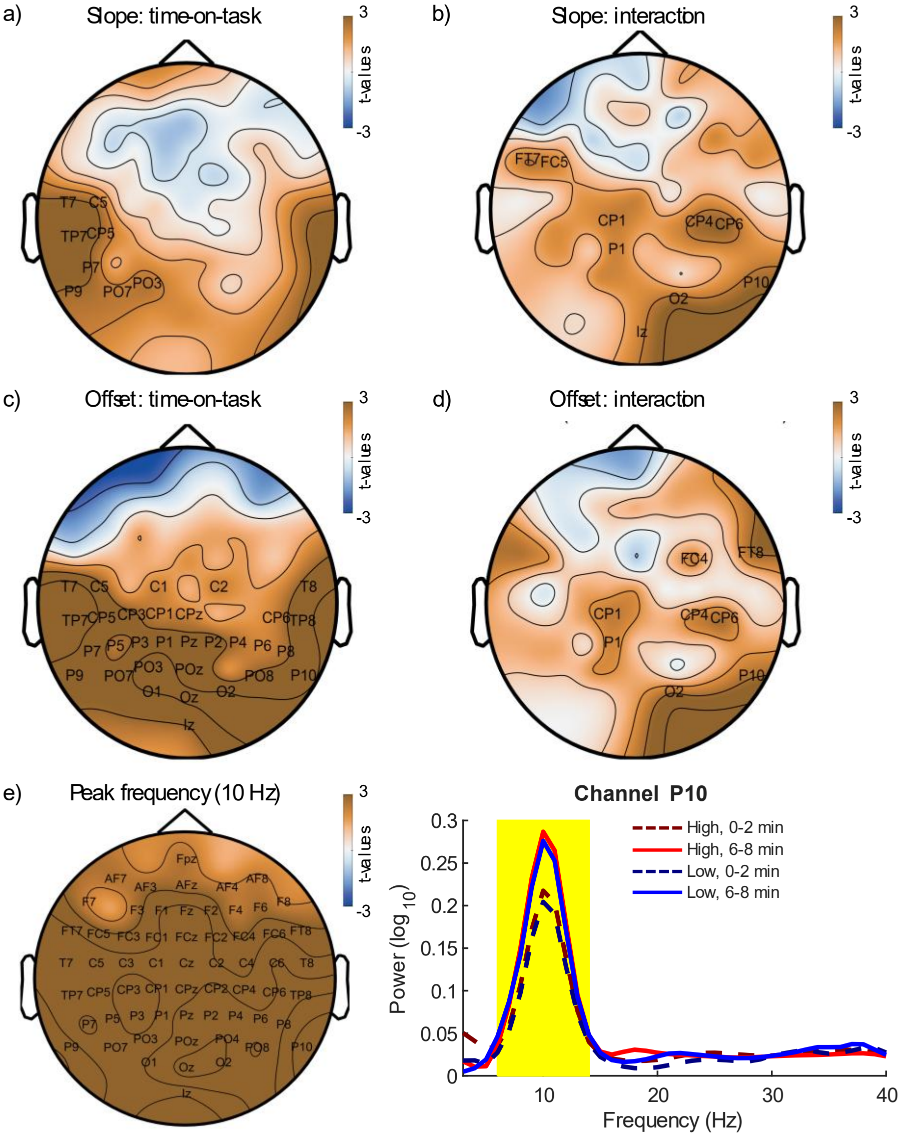


**Supplementary figure 1**. Induced PSD results. a) Scalp topography of the main effect of time-on-task (6-8 min minus 0-2 min) on slope (channel labels are shown for the significant cluster). b) Scalp topography of the interaction of time-on-task with perceptual load (time-on-task effect on high load minus time-on-task effect on low load) on slope. Labels are shown for the significant cluster of channels. c) Scalp topography of the main effect of time-on-task (6-8 min minus 0-2 min) on offset (channel labels are shown for the significant cluster). d) Scalp topography of the interaction of time-on-task with perceptual load (time-on-task effect on high load minus time-on-task effect on low load) on offset. Labels are shown for the significant cluster of channels. e) Scalp topography of time-on-task effect of the periodic component at 10 Hz (6-8 min minus 0-2 min), channel labels are shown for significant cluster of channels, and average power spectrum over participants at channel P10 across time-on-task and perceptual load conditions. Yellow shadow represents the significant frequency range (6-14 Hz) for time-on-task effect.

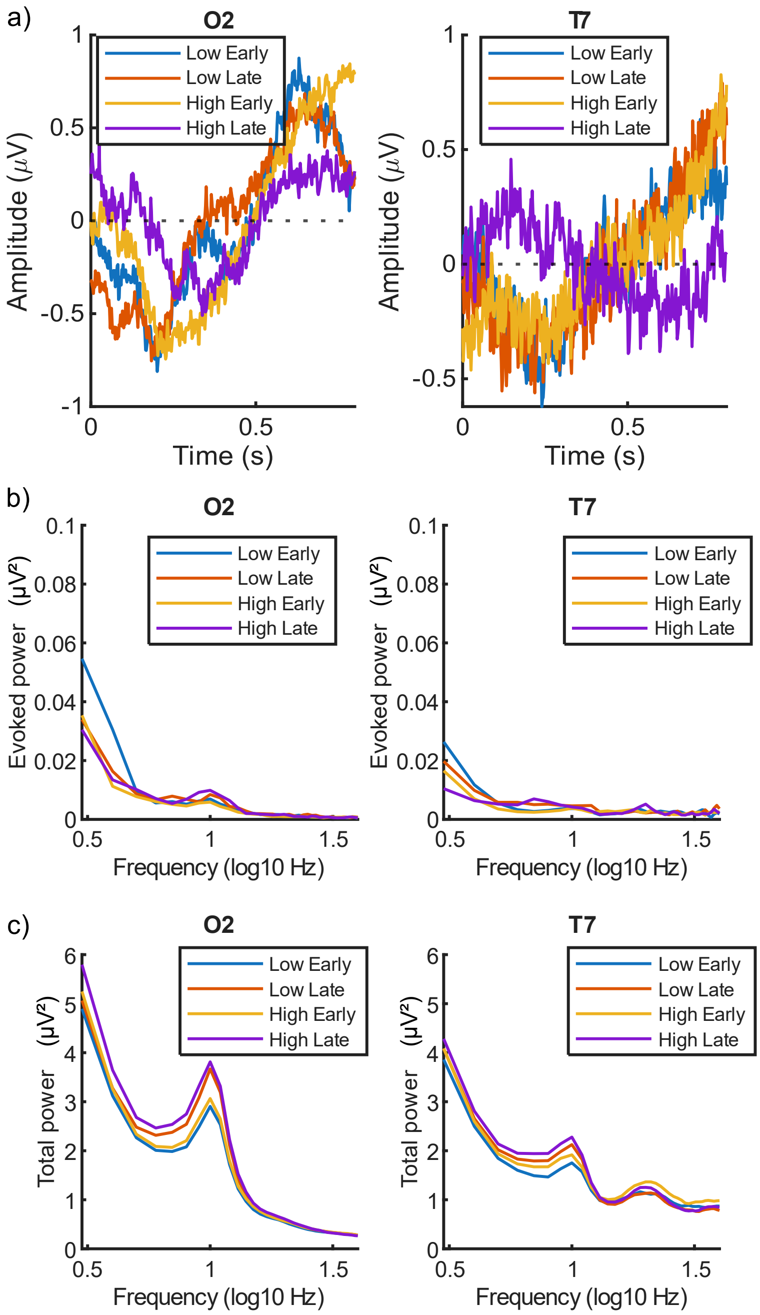


**Supplementary figure 2**. Grand-average activity at electrodes O2 and T7 as a function of perceptual load (low, high) and time-on-task (early: 0–2 min; late: 6–8 min). a) Event-related potentials (ERPs), time-locked to scene onset. b) Evoked power spectral density (PSD), computed from the trial-averaged signal. c) Total PSD, computed from single trials and then averaged, capturing both phase-locked and non-phase-locked activity.
